# A theoretical mechanism for ocular emmetropization

**DOI:** 10.1101/2025.06.01.657224

**Authors:** David Crewther, Nina Riddell, Sheila Crewther

## Abstract

Currently, the human myopia epidemic is estimated to affect nearly 3 billion persons, yet experimental refractive error research is still hampered by the lack of a plausible theoretical mechanism explaining how the eye detects defocus and grows to minimise error (emmetropization). Applied lens defocus, either positive or negative, to animals including human, induces rapid changes in both refraction and axial length. Such changes have been linked to the rate of transfer of fluid from the vitreous chamber across the Retinal Pigment Epithelium (RPE). However, the theoretical basis of sensing the sign of defocus still eludes, as does the physiological operationalisation of this defocus signal. We propose that the signal for the sign of defocus is contained within the pattern of temporal modulation of light on the retina during the myriad saccadic eye movements performed every minute. This pattern is generated by a combination of the ocular point spread function for the defocused eye, narrow cone photoreceptor acceptance of light and the modification of photoreceptor directionality by the retinal shear that accompanies saccadic eye movements. Thus, under conditions of defocus, performing a saccade across a simple visual grating produces an out of focus dynamic stimulus pattern on the retina which is sawtooth-like, the profile being dependent on the sign of defocus. Positive lens defocus induces a fast-OFF/slow-ON sawtooth-like luminance modulation transfer function, while negative lens defocus results in a fast-ON/slow-OFF temporal pattern. Such patterns produce relative incremental or decremental changes in the trans-epithelial electrical potential and consequent changes in fluid absorption across the RPE. Thus, rearing with positive lens defocus is associated with an increase in trans-epithelial potential, increasing trans-epithelial absorption and decreasing vitreous chamber size, while rearing with a negative lens decreases the trans-epithelial electrical potential and decreases RPE fluid absorption, increasing eyeball size. In both cases, retinal images become more in-focus, and the refractive error tends to zero. The resultant is emmetropization, modifying eyeball size in response to applied defocus in a stable negative feedback fashion.

## Introduction

Clear focussed vision occurs naturally in sighted animal species, an ecological necessity. However, in human, myopia (short-sightedness) has increased in prevalence to 36% in children and adolescents (Liang et al., 2024) globally, associated with lifestyle and demographic changes (Saw et al., 2001), especially education (Fan et al., 2014). Animal models of induced refractive error became established in the 1970s when monocular deprivation was shown to result in axial elongation in monkeys (Wiesel & Raviola, 1977) and extreme refractive myopia in chicks (Wallman et al., 1978). Subsequently, the eyes of chicks reared with plus or minus lenses were shown to respond in a differential manner to the sign of defocus. Rearing with a positive lens for a few days results, on lens removal, in hyperopia and a shorter eye, while rearing a with a negative lens results in myopia and a longer eye (Schaeffel et al., 1988). Experimental compensation to defocus is ubiquitous across animal species studied, including macaque monkey(Smith et al., 2014), chick(Wallman et al., 1978), mouse (Barathi et al., 2008), tree shrew (Metlapally & McBrien, 2008), guinea pig (Howlett & McFadden, 2009), squid (Turnbull et al., 2015), marmoset (Troilo et al., 2000) and human (Delshad et al., 2020). Luminance flicker during rearing suppresses myopia development in a frequency dependent manner (Crewther, Barutchu, et al., 2006 ; Schwahn & Schaeffel, 1997), while moving sawtooth luminance modulation, resulting in perceptual brightening or darkening in human (Cavanagh & Anstis, 1986), disrupts emmetropization in chicks in a defocus sign-dependent manner (Crewther & Crewther, 2002).

Molecular science has yielded discoveries of molecules whose expression is differentially related to the response to rearing animals with signed defocus. The transcription factor ZENK (zif268; Egr-1), found in glucagonergic amacrine cells in chick, was the first molecule (Bitzer & Schaeffel, 2006; Fischer et al., 1999) of many genes/proteins, to show differential expression (National Academies of Sciences & Medicine, 2024). Gene pathway analyses have also shown significant structural, metabolic, and immune processes in defocus sign-dependent responses (Riddell & Crewther, 2017; Riddell, Giummarra, et al., 2016).

Retinal pharmacological intervention has impacted on emmetropization, with several attempts to manipulate neurotransmitter function in emmetropization studies. The two isomers of aminoadipic acid interfere with compensation in a way that points to the importance of the balance of retinal ON and OFF responses(Crewther & Crewther, 2003). Similarly, blockers of ion exchangers and cotransporters produce defocus sign - dependent disruption to emmetropization (Crewther et al., 2008). Homatropine (muscarinic acetylcholine receptor antagonist) used in cycloplegic eyedrops for myopia therapy, interferes with choroidal response in defocus sensitive ways (Sander et al., 2018). However, to date, no drug intervention study has hypothesized convincingly how the retina detects defocus.

The role of temporal processing in emmetropization was introduced into the literature by Crewther (2000) and Crewther, Liang, et al. (2006) who proposed that a model based on the processing of spatio-temporal contrast rather than a purely spatial examination of blur was necessary. Rucci and Victor (2018), in supporting this view, posed the question of whether eye movements can contribute to emmetropization. They proposed that small eye movements continually transform the viewed scene into temporal modulations on the retina, without providing direct evidence of how this might affect refractive compensation.

The aim of this manuscript is to develop a theoretical mechanism that explains the process of emmetropization – whereby eye growth compensates to defocus so that the retina becomes optically conjugate with its visual environment (Hodos & Erichsen, 1990). Two questions emerge:- how does the retina signal the sign of defocus, and what is the means by which eye growth rate responds to this signal of defocus?

### Current (incomplete) theories of refractive compensation

There are three commonly referenced incomplete theories related to emmetropization:

1. ***Choroid moving the retina into focus:*** During the induction of form deprivation myopia (FDM) through occlusion, the choroid becomes thinner (Akyol et al., 1996; Liang et al., 2004; Shih et al., 1993; Wallman et al., 1995; Wallman & Winawer, 2004), and during recovery from FDM (the eye is defocused upon occluder removal - highly myopic), the choroidal lymphatics swell greatly, exceeding normal retinal thickness) (Junghans et al., 1999; Liang et al., 2004). Wallman et al. (1995) proposed that the choroid pushes the retina toward a better state of focus. However, in the period of recovery from the time of maximal choroidal thickness (>72 hr after occluder removal with Rx = -8D), the choroid is getting thinner(Liang et al., 2004) while refraction still heads towards zero (Crewther, Liang, et al., 2006). Furthermore, the choroidal push theory does not propose a mechanism for detecting the sign of defocus. Modern optical coherence techniques show extremely rapid responses in axial length, within 2 minutes to applied defocus of either sign in human (Delshad et al., 2020) and there are reports of rapid changes in choroidal thickness in both animals and humans (Wang et al., 2016; Zhu et al., 2005).
2. ***Longitudinal chromatic aberration:*** The focal power of the optics of the chick eye is about 3 dioptres less for red light than for blue light (Rohrer et al., 1992), while in human, this longitudinal chromatic aberration (LCA) for infants is about 1.6 D (Wang et al., 2008). However, chicks emmetropize to applied defocus in monochromatic light (Rohrer et al., 1992; Schaeffel & Howland, 1991; Wildsoet et al., 1993). Normal chicks raised in monochromatic light become slightly more myopic in red light and slightly more hyperopic in blue light (Seidemann & Schaeffel, 2002) compared with white light controls (perhaps simply emmetropizing to different focal planes). Thus, while the pattern of chromatic fringes around object borders might give a learned response for the sign of defocus in humans (Fincham, 1951), it is difficult to explain how a hatchling chick could immediately begin to emmetropize. Also, rearing chicks in red-biased light for four weeks is reported to cause progressive myopia, while rearing in blue-biased light caused progressive hyperopia (Foulds et al., 2013). Resolution of these conflicting findings between monochromatic and spectrally biased rearing is currently lacking.
3. ***Change in rate of fluid transfer across the RPE as the primary driver of ocular growth compensation:*** The RPE is a tight junction-linked monolayer of polarized cells forming apical and basal membranes yielding an apical-positive TEP (trans-epithelial potential) of 4-15 mV (Gallemore et al., 1997; Reichhart & Strauss, 2014; Strauss, 2005). Fluid absorption (in the range 2-11 µl/cm^2^.hr) in the vitreous to choroidal sense is largely controlled by ion exchangers and co-transporters (NaK-ATPase, NKCC1) found on the apical membrane and chloride and bicarbonate channels on the basal membrane. Fluid absorption is related to the magnitude of the trans-epithelial potential (TEP) (Edelman & Miller, 1991; Rymer et al., 2001). The RIDE (Retinal Ion Driven Efflux) model (Crewther, 2000; Crewther, Liang, et al., 2006; Liang et al., 2004) proposed the rate of transfer of fluid across the RPE between the vitreous chamber and the choroidal lymphatic sinusoids (Junghans et al., 1999) as the vector for implementing defocus-induced change in ocular growth. However, while suggesting a mechanism for effecting emmetropization, and allowing biometric ocular changes with the rapidity of ionic changes in the RPE, it does not solve the puzzle of sensing the sign of defocus.

### A theoretical mechanism of emmetropization

The structure of the proposed theoretical mechanism (see Figure 1) uses the wave-guiding properties of photoreceptors, as characterized by the Stiles-Crawford (SC-1) effect (Stiles & Crawford, 1933) and changes in photoreceptor directionality during saccades which exert shear stress on the retina (Richards, 1969). It also relies on known electrical and physiological properties of the retinal pigment epithelium (RPE) (Crewther, 2000; Edelman & Miller, 1991; Gallemore et al., 1997).

**Figure 1.**
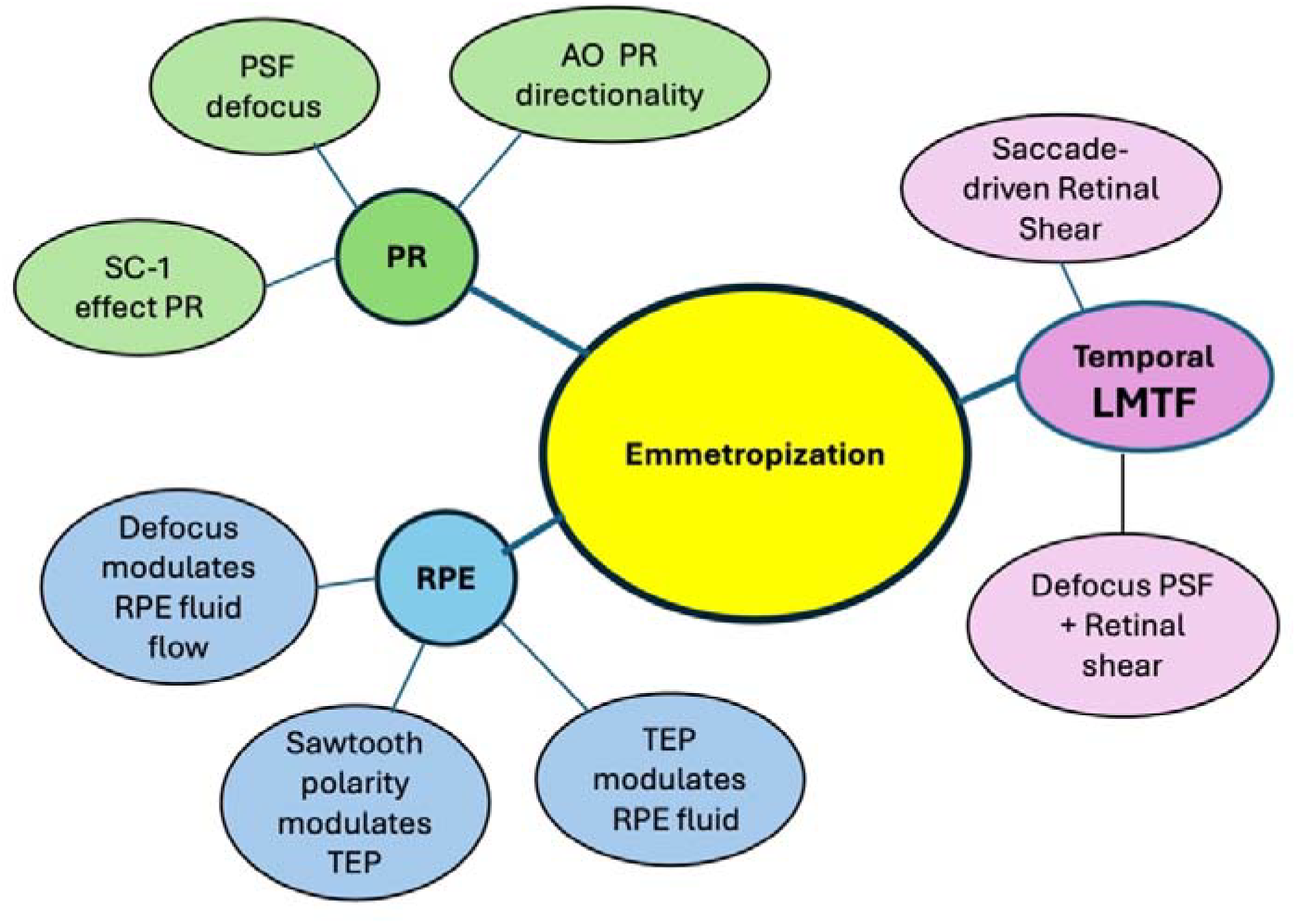
Mind map of the theoretical mechanism of emmetropization. Abbreviations: PR photoreceptor; SC-1 Stiles-Crawford effect of the first kind; PSF point spread function; LMTF luminance modulation transfer function; AO adaptive optics; TEP trans-epithelial potential; RPE retinal pigment epithelium; RIDE Retinal Ion Driven Efflux model.

#### Photoreceptor directionality

Stiles and Crawford (1933) noted that rays entering the centre of the pupil are perceptually brighter than those entering at the periphery of the pupil (SC-1). Indeed a variation in the angle of incidence on a photoreceptor from 0° to 5° requires an increase of about 70% in off-axis intensity in order to give perceptual brightness equality (Pask & Stacey, 1998). The Stiles-Crawford function also differs between photoreceptor types, with retinal rods showing a broader SC-1 peak than do cones. Hence, the effect of eccentric pointing for rods is likely to be less than for cones and hence less relevant for emmetropization.

Fincham (1951) incorporated knowledge of the Stiles-Crawford effect in studying the stimulus to accommodation under defocus of either sign, suggesting that the eye may respond to the vergence signal by using sign-dependant brightness differences across effective blur circles (ePSF) due to the photoreceptor eccentricity away from the fovea. He postulated that there would be a peripheral retinal signal for defocus sign of the blur circle on the retina (believing the photoreceptors pointed to the centre of the eye). Fincham speculated that small eye movements across the fovea would create a defocus sensitive signature, though his theory did not extend to translating such a signal to eye growth changes. However, the advent of adaptive optics (AO) techniques applied to individual human cones showed that that their directionalities converged to the centre of the pupil (or slightly nasally) (Roorda & Williams, 2002). The corrected theory, taking account of photoreceptor intrinsic phototropism, lost the means to sense defocus sign (Kruger et al., 2001).

#### Photoreceptor decentration via saccadic retinal shear

Retinal tissue is soft by comparison with choroid (10x stiffer) and sclera (500x stiffer) (Aboulatta et al., 2021; Qu et al., 2018). Every time the eye makes a saccadic eye movement, there is rapid rotational acceleration, generating a shear force affecting photoreceptor directionality (sensitivity to light coming from different directions). That shear force is transferred from sclera to choroid and then to the retina. This alters the directionality of the photoreceptors, consistently opposed to the direction of scleral movement of the saccade. As early as 1969, this “retinal shear” was measured through the shift in the peak of the Stiles-Crawford function during saccades (Richards, 1969). For human, the average shift in the Stiles-Crawford effect is about 0.6 mm in the pupil plane measured at a time 40ms after the start of a saccade of amplitude 5°. This corresponds to about a 3° angular tilt for the retinal photoreceptors. This may underestimate the maximum photoreceptor tilt as most 5° saccades are almost complete by 40 msec (Robinson, 2022b), the eye coming to a stop, with the remaining bend in photoreceptors dependent on the shear relaxation time.

Large saccades (>10°) across a grating of 1 cpd produce temporal frequencies higher than photoreceptors can resolve. For example, main sequence saccades (Robinson, 2022b), show peak velocities of 350°/s for saccades of 10° in amplitude. Thus, the pattern projected onto the retina would possess a frequency of nearly 200Hz, whereas cones are temporally limited to 60-90 Hz (Hamer & Tyler, 1992; Kelly, 1971). Saccades of amplitude < 2-3° are sufficiently small to provide temporal modulation of photoreceptors within physiological range, under such a scenario (Rucci & Victor, 2018). There is current active discussion in the literature about whether vision proceeds by a series of saccade driven snapshots morphed together or through the temporal sequence of events experienced through the retina (Gur, 2024; Rucci et al., 2025), however the same question may be raised with the question of emmetropization - is it a matter of adjustment of spatial blur or a feature of the response to the temporal code of illumination on the retina (Crewther, 2000; Crewther, Liang, et al., 2006; Rucci & Victor, 2018).

## Methods

The standard reduced emmetropic eye is sufficient for illustrating the mechanism of emmetropization. Hence, we can avoid discussing lens contributions and even aberration types. The geometrical optics blur circle for defocus of *Δ*E dioptres on the retina of an eye with pupil diameter D metres has a diameter b metres given by (Tunnacliffe, 2021)

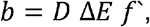

where *f*’ is the posterior focal length of the eye.

In order to take account of photoreceptor directionality, the Stiles-Crawford function of the first kind (SC1) is typically modelled as a Gaussian function:

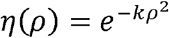

where *ρ* is the distance from the centre of the pupil measured in the pupil plane and *k* is a scaling constant that determines the width of the sensitivity drop-off (typically *k* ≈ .05 − .07 *mm*^-2^ for humans).

Richards (1969) quantified a shift in the peak of the SC-1 during saccadic eye movements. He showed that in the pupil plane, the photoreceptor alignment was shifted by 0.6 mm. This equates to about 3°. Richards employed 5° amplitude saccades, making measurements 40 ms after the previous saccade had started (Richards, 1969). However, most saccades of this amplitude have a duration of 30-40ms (Robinson, 2022a). Richards states that 15 ms after the eye stops moving the retina still trails the sclera, and the photoreceptors would be in a state of shear relaxation. Thus, the maximum decentration of photoreceptors would likely be larger.

The retinal shear effect depends on the rotational acceleration of the eye after the start of the eye movement and on the shear modulus of the retina as well as the viscoelastic properties of the vitreous humor on the inner limiting membrane at the interface of the retina and vitreous humor. The shear relaxation time time for retinal vibration has been recently measured using optical coherence elastography (OCE) techniques (He et al., 2021; Zhang et al., 2020; Zhang et al., 2018). Typically, two time constants are evaluated the fast recovery time *τ*1, of retina is ≈100ms. Hence, if the eye performs 3 saccades per ssecond, the photoreceptors are decentred, consistently against the direction of motion of the retina for roughly 30% of the time. Hence, in a fashion like Fincham and Kruger, we can establish the effective point spread function (ePSF) for a defocused eye that are differentiated by the sign of defocus. Adding decentration of photoreceptors shifts the peak of the PSF, now biased to one side or the other, depending on the sign of defocus (as recognised by Fincham (1951)). For photoreceptors decentred by *r*_0_ in the pupil plane, the SC function is is then: 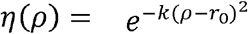. The next step is to calculate the effective luminance modulation transfer (eLMTF) tasking account of the defocused, decentred (bent) photoreceptors. A LabVIEW (ni.com) program was designed which employed pupil diameter, spread of the SC-1 function and the degree of decentration of cone directionality as input variables to generate eLMTF for plus and minus lens defocus. The convolution of the temporal square wave and the decentred, defocused ePSF yields a temporal modulation (eLMTF), as shown in Fig. 2.

**Figure 2.**
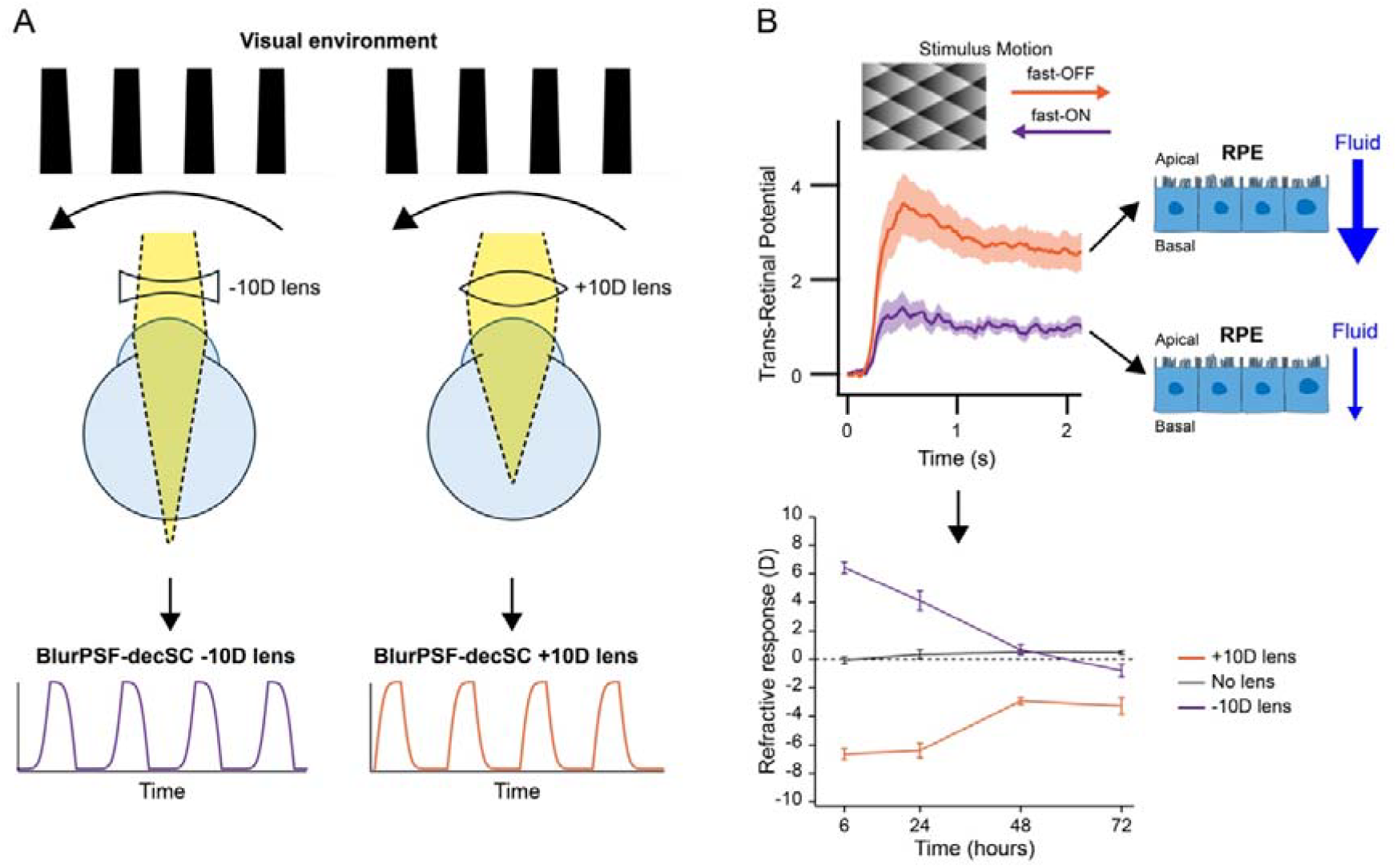
**A** The biometrics of emmetropization with rapid response to defocus. Rapid eye movement across a grating stimulus produces different sawtooth luminance modulation transfer function on the retina depending on the sign of defocus. **B** Trans-retinal potential effects of sawtooth stimulation with shaded diamonds. A brightening stimulus (fast-OFF slow-ON) results in an increased trans-retinal potential compared with the same stimulus moving the reverse direction creating a fast-ON slow-OFF pattern. A fast-OFF temporal pattern increases fluid efflux while fast-ON reduces fluid efflux.

## Results

### Temporal processing of a blurred visual stimulus during a saccade - the signal of of defocus sign

Even when visually fixating, we execute 1 - 2 small (<1°) fixational eye movement (microsaccades) per sec of which we are not aware (Martinez-Conde et al., 2004). Also, we make multiple small saccades per second when reading, of amplitude 1-3° (Rayner, 1998). Thus, the period of visual stimulation with transiently decentred photoreceptors represents a considerable fraction of total time. This is important, as the signal for defocus and the means of controlling eye growth are closely coupled as is discussed below. While early attempts (Fincham, 1951; Kruger et al., 2001) to solve the puzzle of signed defocus considered the static point-spread function (PSF), we now consider the temporal pattern of visual stimulation on the retina during an eye movement. Consider a saccadic eye movement from a target on the right to one on the left across a static stimulus – say, a square-wave grating (see Fig 2A). The eye makes a saccade with rapid acceleration of the eye. For an in-focus retina, a patch of the retina would receive a temporal square wave of illumination. The effect of blur plus the decentred Stiles Crawford effect on the eLMTF is shown for positive and negative lens defocus ion Fig. 2A

When the Stiles-Crawford function of the eye (Gaussian in shape) is taken into consideration, the distribution of photo-absorption by the retina becomes an intersection of the blur circle and the Gaussian. Adding decentration of photoreceptors shifts the peak of the PSF, now biased to one side or the other, depending on the sign of defocus (as recognised by Fincham (1951)). The LabVIEW program employed pupil diameter, spread of the SC-1 function and the degree of decentration of cone directionality as input variables to generate eLMTF for plus and minus lens defocus. The program calculates the temporal waveform when defocus, the application of a centred SC-1 function and the effect of decentration of the cone photoreceptors are considered. The convolution of the temporal square wave and the decentred, defocused ePSF yields a sawtooth-like luminance modulation transfer function (eLMTF) with either fast-OFF (for positive lens defocus) or fast-ON (for negative lens defocus) – see Fig. 2.

### Effects of Sawtooth Pattern Stimulation on the DC-Electroretinogram

The ubiquity across the animal world of clear focused vision argues for a common mechanism to detect defocus and its sign. Thus, Riddell, Hugrass, et al. (2016) used an isolated toad eyecup, a stable physiological model, to investigate the effect of sawtooth temporal profiles on the trans retinal potential (TRP - measured between vitreous chamber and sclera). Computer-generated (VPixx.com) dynamic images were projected onto the retina. Drifting sawtooth gratings (including shaded diamond patterns) showed a significant difference in the TRP (see Fig 2B) between the fast-OFF and the fast-ON patterns (Riddell, Hugrass, et al., 2016). This difference was maintained at different drift rates yielding a temporal stimulation frequency of at least 20Hz.

A relation between TEP or TRP and fluid absorption across the RPE is accepted across species – in frog/toad (Adorante & Miller, 1990; Edelman et al., 1994; Hughes et al., 1988; Miller et al., 1982), in cow (Edelman & Miller, 1991; Peterson et al., 1997; Rymer et al., 2001),in dog(Tsuboi, 1987), in monkey (Tsuboi & Pederson, 1988), in chick (Li et al., 1994; Wolfensberger et al., 1999) and human (Adijanto et al., 2009; Maminishkis et al., 2002). These show, in general, a positive relation between TRP (or TEP) and fluid flow.

## Discussion

In a recent review, Schaeffel and Swiatczak (2024) proposed that emmetropization is due to an autofocus system with negative feedback. We agree, and propose that such a defocus sensitive system is provided by the temporal pattern of illumination on the retina during the many small eye movements that we perform without effort or often, without knowledge. When an eye changes between a state of hyperopic defocus and myopic defocus, the eLMTF during saccades changes accordingly, rendering stable negative feedback through the change in trans-epithelial potential and associated fluid absorption rate across the RPE. The speed of refractive and biometric change afforded by this theory conforms with the measured rapidity in animals and human, likely due to the speed with which electrical and fluid changes in the RPE occur.

Because the photoreceptors bend under saccadic acceleration – always against the direction of motion of the sclera, a realistic difference is induced between plus and minus lens power on the defocused temporal eLMTF patterns played out on the retina. When the temporal modulation of the light pattern of a grating on the retina during eye movements becomes sawtooth-like with polarity and transepithelial potential differences according to sign of defocus, the mechanism mimics a machine, increasing or decreasing pump rate of fluid across the RPE, with an integrated sensor/effector system for changing the vitreous chamber shape in response to defocus. Riddell, Hugrass, et al. (2016) investigated the neural basis of these phenomena. The addition of APB (known to eliminate the electroretinogram ON-response) abolished the peak at drift offset but increased the DC difference between the effects of fast-OFF and fast-ON stimulation, implicating the RPE as the main neural source of the DC effect.

Do all stimulus motions across the retina contribute to growth signals? Without retinal shear associated with rapid rotational acceleration of the eye, there is no consistent tendency to decentre the directionality of photoreceptors in a direction against the motion of the sclera. Hence, unless the photoreceptors are bent, directed away from their stationary direction (centre of the pupil) refractive compensation will not occur.

The mechanism has been illustrated with very simple conditions – a reduced eye and the effects of making a saccade across a simple square wave grating. The generalization to complex visual patterns is straightforward. Just as with Fourier expansions, which use sine and cosine functions with variable frequency there are complete basis sets of square waves such as the Walsh functions with its associated Walsh-Hadamard Transform. In transforming the spatial environment to temporal modulation of light on a locus of the retina, accompanying a saccadic eye-movement, the higher spatial frequencies of the general function turn into higher temporal frequencies of the eLMTF. This should not interfere with the electrophysiological effects as we know the differential effect of sawtooth patterns on the TRP certainly span 2-20 Hz at least (Riddell, Hugrass, et al., 2016).

In terms of the other theories of refractive compensation referenced above, some unification is possible. The rapid increases or decreases of choroidal thickness associated with lens defocus are consistent with fluid transfer into or from the lymphatic sinusoids (Crewther, Liang, et al., 2006) under control of RPE, in turn reflecting the temporal pattern of stimulation of the cone photoreceptors. In terms of theories based on chromatic aberration, the current theoretical mechanism would drive emmetropization to its spectral focal endpoint, were there monochromatic light used. In terms of differences in the SC functions between different cone types, one would expect the same anatomical structure of L and M cones, however, little is known about the individual S cone directionality through reflectometry.

## Conclusion

Saccadic eye-movements cause retinal shear that consequentially affect light directionality and intensity in photoreceptors. When coupled with positive lens defocus, moving the eyes across a patterned visual environment results in a fast-OFF/slow-ON temporal modulation of retinal illuminance causing an increase in trans-retinal potential and accompanying increase in fluid outflow, resulting in relative reduction in vitreous chamber depth. With negative lens defocus, the temporal pattern is biased toward fast-ON/ slow-OFF temporal modulation, causing a decrease in trans retinal potential and a decrease in fluid outflow, resulting in a relative increase in vitreous chamber depth.

Hence, emmetropization – the process of bringing the eye into focus no matter the sign of blur, is a natural active automatic photoreceptor mechanism. A positive lens increases outflow and causes relatively shorter vitreous chamber. Conversely, negative defocus decreases outflow and causes a relatively longer vitreous chamber.

## Acknowledgments

Commercial relationships: none.

